# Explicit memory representations in decisions from experience

**DOI:** 10.1101/2025.10.27.684917

**Authors:** Alice Mason, Marcus Lindskog, Ralph Hertwig, Dirk U. Wulff

## Abstract

Reinforcement learning (RL) models explain how people adapt behavior through incremental value updates but assume that individual experiences are not explicitly stored or retrieved. Across two experiments (N = 282 and N = 1,818), we tested whether people rely on such explicit memory representations during experience-based choice. Participants sampled outcomes from two lotteries and, in an “ignore” condition, were instructed to disregard specific outcomes before deciding. Ignoring these outcomes substantially altered preferences, suggesting that choices were guided by explicit episodic representations rather than cumulative reinforcement. Frequency judgments revealed generally accurate memory for experienced outcomes, but reduced precision for continuous compared with discrete outcome distributions. These findings challenge purely incremental RL accounts and support theories proposing that human choice integrates flexible episodic memory with reinforcement mechanisms, bridging models of learning, memory, and decision-making.

**Statement of Relevance:** How people learn from experience shapes decisions in many everyday situations, from choosing a stock to invest in to selecting a restaurant or deciding which route to take home. Our experiments show that people do not rely solely on incremental value updates but instead exploit memory representations of past outcomes to guide choices. When asked to ignore certain outcomes, participants adjusted their preferences in line with these representations. These findings reveal that human decision-making also engages flexible use of memory for past experiences, highlighting the importance of memory-based mechanisms in adaptive, real-world choice behaviour.

## Introduction

Reinforcement learning (RL) offers a fundamental framework for how people learn from experience, explaining how outcomes shape future decisions. It has been applied across domains from machine learning and artificial intelligence (Christiano et al., 2017; Silver et al., 2017; Sutton & Barto, 1998) to human decision-making in risky choice and strategic games (Decker, Otto, Daw, & Hartley, 2016; Gershman & Daw, 2017). A core assumption of RL is that learning occurs incrementally through value updates, without forming explicit memories of individual outcomes. Yet, it remains unclear whether people rely solely on these summary statistics or also maintain richer, detailed representations of past experiences when making decisions.

Empirical studies of risky choice often use repeated samples from monetary lotteries or multi-armed bandit tasks and model behavior using RL frameworks(Palminteri & Lebreton, 2022; Spektor & Wulff, 2021; Weber, Yee, Small, & Petzschner, 2025). While RL models capture many important behavioral patterns (Daw & Doya, 2006), they can perform poorly under conditions of noise or limited data (Gershman & Daw, 2017; Lengyel & Dayan, 2007). These shortcomings suggest that decision-makers may rely on more detailed representations of past experiences than standard RL models assume, motivating the consideration of memory-based mechanisms to better account for choice behavior (Bornstein, Khaw, Shohamy, & Daw, 2017; Gershman & Daw, 2017; Spektor & Wulff, 2024).

An alternative hypothesis is that decision-makers rely on rich memory representations of past experiences. Several influential models in the decision-making literature capture behavior by assuming that individuals sample past outcomes in ways consistent with well-established memory mechanisms. For example, items may be sampled according to similarity to the current context (Dougherty, Gettys, & Ogden, 1999; Gonzalez & Dutt, 2011), recency (Erev, Glozman, & Hertwig, 2008; Plonsky, Teodorescu, & Erev, 2015; Spektor & Wulff, 2024), interference, or saliency (Hotaling, Jarvstad, Donkin, & Ben, 2019), or combinations of these factors. While these models can successfully reproduce choice behavior, little is known about how individuals actually represent individual episodes in memory or what the boundary conditions are for such representations. Most research has not directly assessed or manipulated memory representations, instead relying on computational assumptions (Olschewski, Luckman, Mason, Ludvig, & Konstantinidis, 2023; Spektor & Wulff, 2024; Wulff & Pachur, 2016), with a few exceptions (Mason, Madan, Simonsen, Spetch, & Ludvig, 2022).

Here, we directly test whether people form explicit representations of outcomes and options during risky choice tasks, asking how the sequential experience of information is stored and used to guide subsequent decisions.

### The current study

We used a between-subjects manipulation to test how individuals represent sequential information during a decisions from experience task (Hertwig, Barron, Weber, & Erev, 2004; Hertwig & Wulff, 2022; Olschewski et al., 2023; Wulff, Mergenthaler-Canseco, & Hertwig, 2018). This task allowed participants to learn about two choice options by sampling from them. Participants in one group, but not the other, were instructed to ignore specific outcomes before making a choice. These outcomes occurred with different frequencies across the two choice options, so ignoring them systematically altered the relative attractiveness of the options. Importantly, when the to-be-ignored outcomes are excluded, the task reduces to well-studied problems in the decisions-from-experience literature (Wulff et al., 2018), providing a strong baseline for predicting choice patterns. If participants form explicit memory-based representations of the sampled outcomes, their preferences should shift in line with the change in attractiveness and past preference patterns. By contrast, if participants integrate outcomes incrementally, as they are encountered and with no individual experiences being stored, the instruction to disregard particular outcomes should have little impact on choice. In addition to choice, we assessed participants’ judgments of the raw frequencies of outcomes they have experienced, which can only be accurate to the extent that explicit memory-based representations have been formed.

Experiment 1 used simple lotteries with a small number of discrete outcomes, a design that may facilitate explicit encoding. Experiment 2 both replicated this problem and introduced choices with continuous outcome distributions, allowing us to test whether participants maintain explicit, memory-based representations even in more complex environments.

In addition to the ignore manipulation, we varied the instructions by informing half of the participants only after sampling that they would later make a choice between the options. This manipulation was designed to test whether the formation of explicit memory depends on the goals implied by the task structure (Hogarth & Einhorn, 1992; Lindskog, Winman, & Juslin, 2013). The instruction-timing manipulation had no effect on choices or frequency judgments, so we focus on the findings of the ignore instruction. Finally, because the task involves numeric reasoning, we also examined whether participants’ numeracy, measured via a standard test, moderated these effects.

## Methods

### Participants

Participants were recruited through the Prolific Academic web service (https://prolific.ac/). The study lasted approximately 10 minutes. Participants were compensated with £0.75 for their participation, plus a decision-contingent bonus of £0.04 on average (Experiment 1) and £0.12 (Experiment 2). In Experiment 1, 282 people (68% female) participated, aged 19 to 81, with a mean age of 37 (SD = 12.5). In Experiment 2, a total of 1,818 participants (52% female), aged 18 to 75, took part, with a mean age of 35.8 (SD = 12.6). Of these, 961 and 857 participants were assigned to the discrete and continuous conditions, respectively.

### Procedure

We conducted two experiments involving the sampling paradigm (Hertwig et al., 2004; Wulff et al., 2018), a standard task in the literature on decisions from experience. In this task, participants were presented with two boxes, each representing a separate monetary lottery. To learn about lotteries, individuals were allowed to sample exactly 24 times from each of the lotteries in whatever order they wanted. Individuals were instructed to sample until both options became inactive, and they were unaware of the exact number of samples permitted for each lottery. To eliminate noise associated with sampling error, the samples were drawn randomly without replacement from a fixed set of samples for each option (see Table 1). Before participants entered the main task, they practiced the task by sampling a few times from two options with other outcome distributions. In Experiment 1, all participants were presented with discrete outcomes. In Experiment 2, participants were either presented with discrete or continuous outcomes.

**Table 1.**
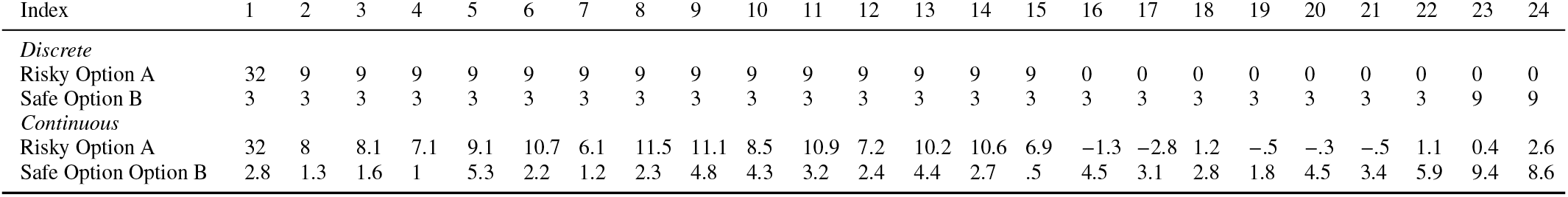
Decision problems.

In both experiments, there were two between-subjects manipulations. The primary manipulation, the ignoreinstruction, encompassed a control group receiving no additional instructions and an experimental group being instructed to ignore particular values and make their choice as if they had not seen these values. For the discrete outcomes, they were instructed to ignore all outcomes of 9, and for the continuous outcomes, to ignore all values between 6 and 15. The secondary instruction manipulation determined whether participants knew from the beginning that they would be making a choice or whether they learned of the choice only after sampling from the two options.

After individuals indicated their choice, they proceeded to a surprise frequency judgment task. Individuals were instructed to judge the raw frequency of each relevant outcome. That is, individuals who were instructed to ignore particular outcomes judged only the frequencies of the remaining three outcomes across the two options. Frequencies were indicated using sliders ranging from 0 to a maximum per-slider value of 50. In Experiment 2, a slightly altered frequency judgment procedure was used. Instead of asking individuals directly for the frequencies of each outcome, we first asked them to judge the total number of samples observed per option and then, in a second step, how these samples were distributed across the observed outcomes. This alternative judgment format was implemented to distinguish between absolute and relative representations of occurrences. In the case of the latter, the two-stage procedure should help provide more accurate frequency judgments.

The options of the decision problem were set up such that ignoring outcomes of 9 (Experiment 1 & 2) or values between 6 and 15 (Experiment 2) in both options (A and B) would achieve two goals. First, it would reduce the relative attractiveness of the risky option (A, see Table 1). Including all outcomes, the risky option would offer a substantially higher average value of £6.58 versus £3.5 and a roughly 60% chance to yield a better outcome than the safe option in a single draw. In Experiment 1 and in the discrete condition in Experiment 2, eliminating outcomes of 9, however, reduces the risky option’s average value dramatically to £3.2, whereas the safer option’s average value is almost unchanged at £3. A similar reasoning holds for the continuous condition in Experiment 2. At the same time, the probability of yielding a better outcome on a single draw by choosing the risky option reduces to 10%. Individuals’ choices were incentivized with a single draw from the chosen lottery and received one-tenth of the drawn outcome as a monetary bonus. The second goal of the ignore instruction was to reduce the choice problem to one that has been studied extensively in the literature on decisions from experience (Wulff et al., 2018). This metamorphosis enabled us to predict where preferences under the ignore manipulation would fall, using data from a recent meta-analysis Wulff et al. (2018).

Finally, individuals completed all four items of the Berlin Numeracy Test (BNC; Cokely, Galesic, Schulz, Ghazal, & Garcia-Retamero, 2012) and answered a few demographic questions.

### Statistical analysis and data

The data were analyzed using R Statistical Software (v4.1.2; R Core Team 2021). The data and corresponding analysis code are available at https://osf.io/d7u5e.

## Results

What is the nature of memory representations in decision from experience? To address this, we first focus on choice, analyzing whether the manipulations and covariates impacted people’s preference for the risky option. Afterwards, we turn to the frequency judgments, analyzing the effect of the manipulations and covariates on the distances between people’s judgments and the true experienced frequencies.

### Risky choice

The results, illustrated in Figure 2, show that choice was heavily impacted by the ignore instruction, suggesting that individuals were able to retrieve stored episodes of outcomes. Specifically, in Experiment 1, individuals instructed to ignore outcomes of 9 had a much lower rate of risky choice 25.3% (95%CI: 18.9 – 33.1) compared to individuals not instructed to ignore outcomes of 9 who preferred the risky option 59% (95%CI: 51.0 – 67.1). In Experiment 2, participants were shown outcomes from either a discrete or a continuous outcome distribution. Similarly to the results for Experiment 1, when the distribution type was discrete, and participants were given the ignore instruction, they chose the risky option 23.6% (95%CI: 20 – 27.6) of the time compared to those in the control condition. The latter opted for the risky option 61.5% (95%CI: 57.1 – 68.5) of the time. Overall, participants who were shown outcomes from the continuous distribution chose the risky option more often than participants shown the discrete outcomes. There was still a strong impact of the instruction to ignore: participants who were asked to ignore chose the risky option 37.7% (95%CI: 32.8 – 42.9) of the time compared to the control condition, which had a higher rate of risky choice, namely, 68.8% (95%CI: 64.7 – 72.7).

**Figure 1.**
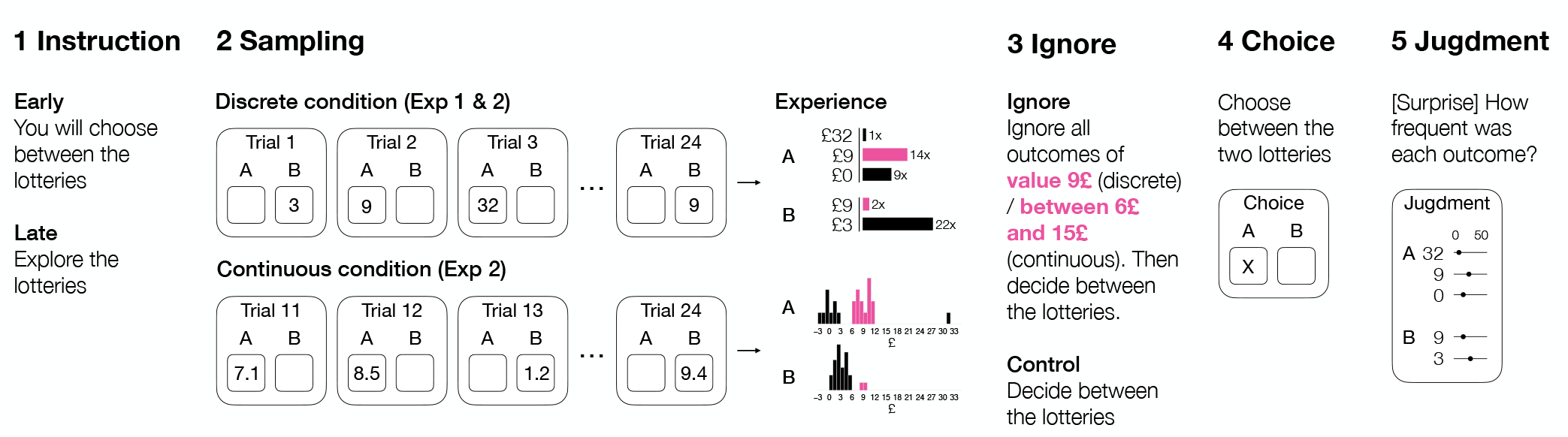
Overview of the study designs. (1) Participants received instructions either before or after the sampling phase on having to decide between two lotteries, (2) sampled outcomes from two either discrete (Experiment 1 and 2) or continuous (experiment 2) choice options across 24 trials, (3) in the ignore condition, were instructed to disregard specific outcomes (£9 for discrete options; £6–£15 for continuous options) before making their choice, (4) made a final choice between the two options, and (5) judged the relative frequency of outcomes in a surprise test.

**Figure 2.**
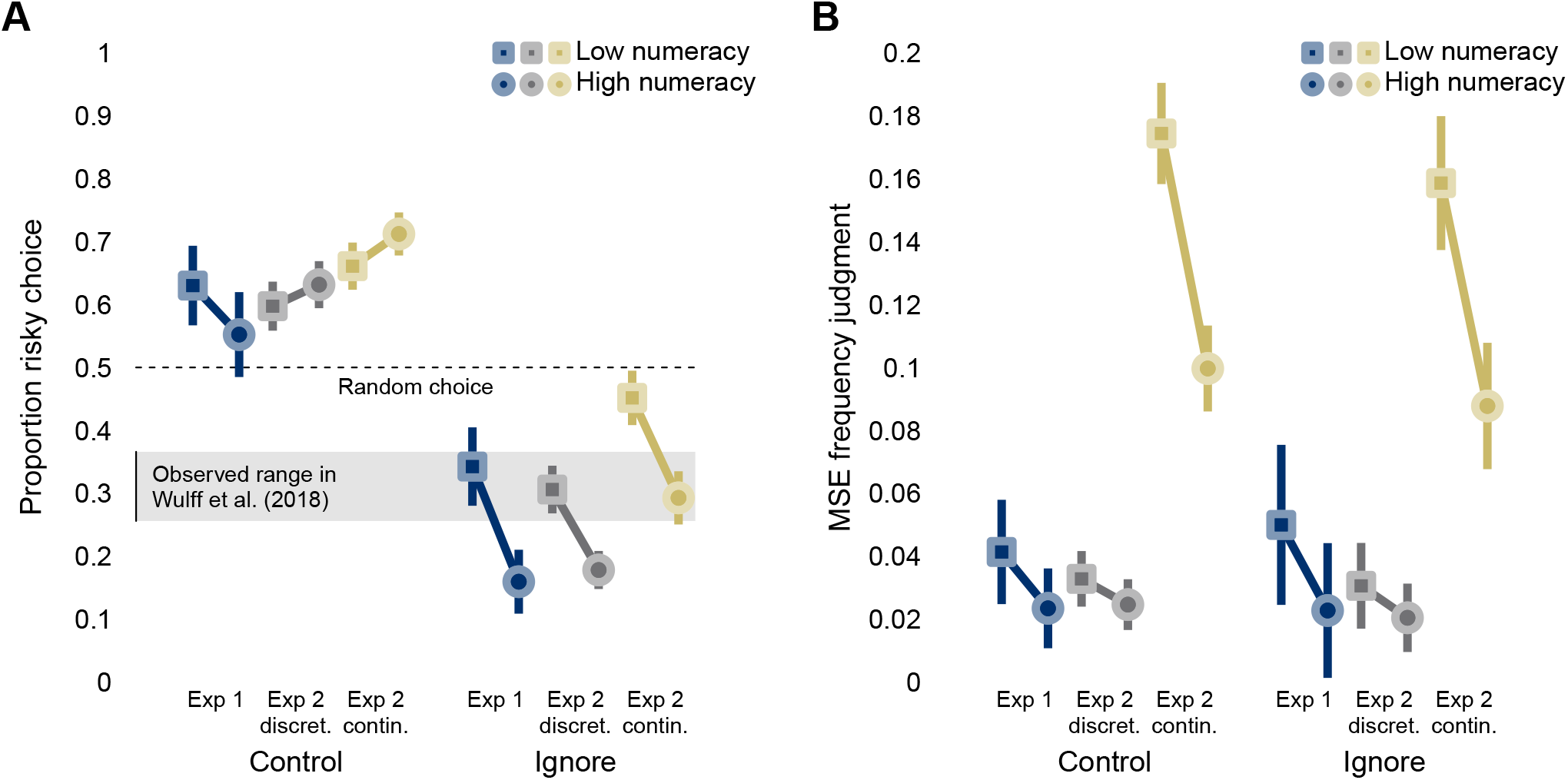
Risky choice and errors in frequency judgments. Panel A shows the proportion of risky choices in the control group and for participants, whereas panel B shows the mean squared error of participants’ probability judgments. The panels show the results from both Experiment 1, where all participants saw a discrete distribution, and Experiment 2, where participants either saw outcomes from a discrete or a continuous distribution. Squares (circles) show the results for above-median (below-median) numeracy scores according to their scores on the Berlin Numeracy Test. The horizontal dashed line in panel A indicates the expected result under random behavior, whereas the shaded area shows the expectation under systematic behavior, which has been derived from the data of 32 studies from a recent meta-analysis (Wulff et al., 2018, see text;).

Crucially, the choice proportions under the ignore instruction in the three data sets (Exp. 1, Exp. 2: discrete and continuous) align well with the choice proportions observed in the past. By following this instruction, the decision problem simplifies into one commonly discussed in the literature on decisions made from experience. Using data from a recent raw-data meta-analysis (Wulff et al., 2018), we analyzed the proportion of risky choices in 32 datasets and found that people in this choice problem preferred the risky option in 31.1% of choices (95%CI: 25.6 – 36.6). This is consistent with the choice proportions we consistently observed in the present data sets.

To examine the effects of each of the key variables, we conducted a binomial logistic regression to predict individual choices (risky option or safe option) from the key experimental manipulations (Ignore Manipulation [Ignore or Control], Instruction Timing [Before or After], Distribution Type [Discrete or Continuous], Numeracy [High or Low]). All variables and their two-way interactions were entered into the model. We evaluated the importance of predictors in terms of the deviance delta between a model with the predictor and a model without. Figure 3 shows that four predictors had statistically significant deviance deltas. First, people who received the ignore instruction were less likely to choose the risky option than those without receiving it (Δ = 264.6, *b* = − 1.51[− 1.70, − 1.32]), amounting to, by far, the largest effect. Second, people who were presented with outcomes from a continuous distribution were more likely to choose the risky option than people who saw discrete outcomes (Δ = 24.6, *b* = 0.48[0.29, 0.67]). Third, people with high numeracy scores were less likely to choose the risky option than people with low numeracy scores (Δ = 8.2, *b* = − 0.27[− 0.46, − 0.09]). Finally, the ignore manipulation interacted with numeracy, with people with high numeracy scores showing a larger difference between the ignore and no instruction than people with low numeracy scores (Δ = 19.7, *b* = −0.85[−1.23, −0.47]).

**Figure 3.**
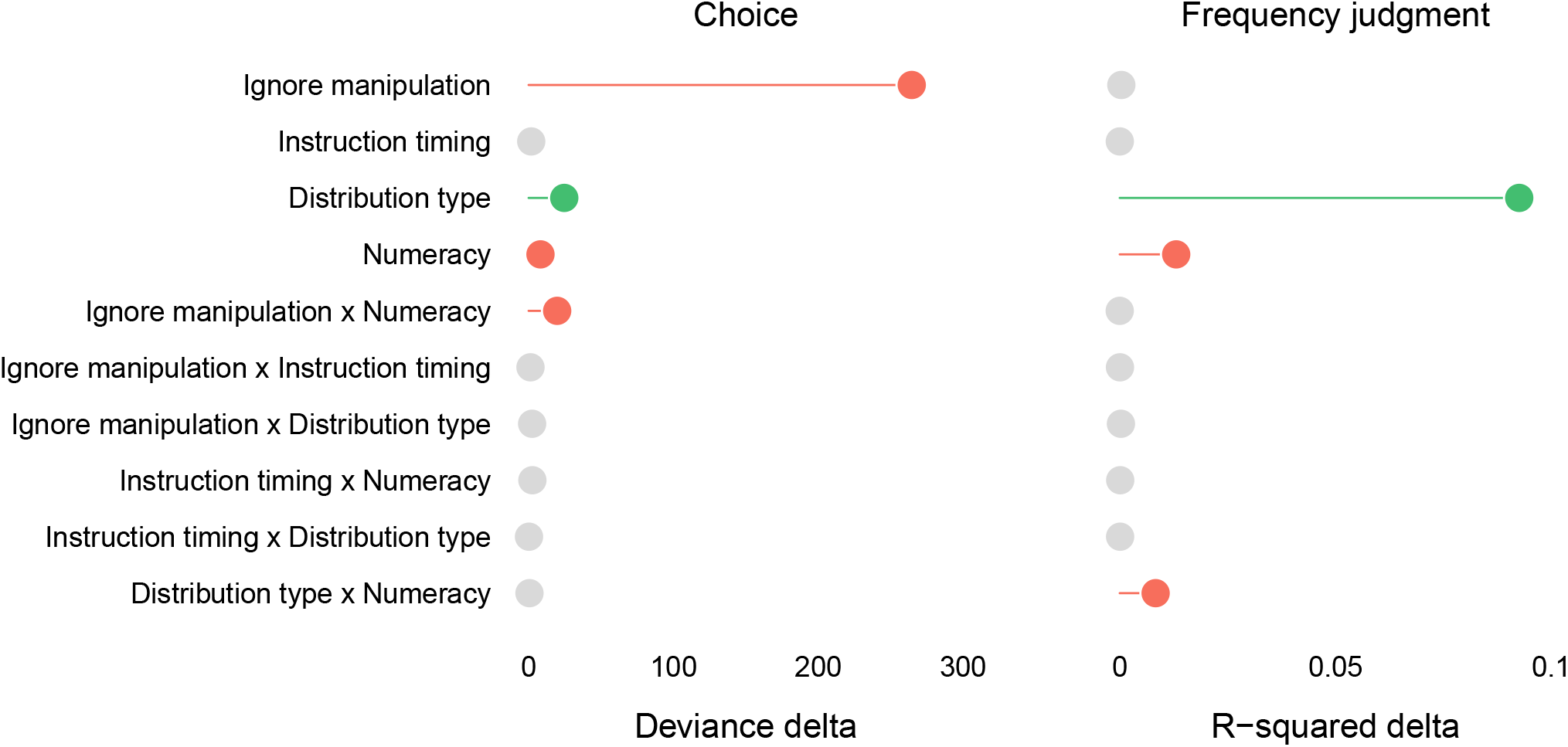
Variable importance for the prediction of risky choice and errors in frequency judgments. The left panel shows the differences in deviance (Deviance delta) between models with and without each predictor predicting risky choice. The right panel shows the differences in explained variance (R-squared delta) between models with and without each predictor predicting the squared error in the frequency judgment task. Colors indicate the direction and significance of the effect, with green implying a significant positive effect, red a significant negative effect, and grey an effect that did not reach significance (according to the model including all predictors).

Overall, the analysis indicates that regardless of when the instruction is given, individuals can effectively adjust their preferences in response to the ignore manipulation, matching choices seen in studies where participants never encountered the outcomes to be ignored. This finding strongly suggests that explicit memory representations of past experiences are present and accessible. Moreover, individuals with strong numeric abilities are more responsive to the ignore manipulation, implying that numeric skills may play a role in accessing or forming memory representations of experiences.

### Frequency judgments

After making a choice, participants were asked, without prior warning, to estimate the relative frequencies of the different outcome options. Figure 2 plots the mean squared error (MSE) of their frequency estimates in the different experimental conditions. Most notably, unlike for choice, the ignore manipulation did not have a significant effect on the accuracy of people’s frequency judgments. In Experiment 1, for people in the control condition, the average estimate error was *MS E* = 0.033(95%*CI*: 0.028 − 0.038), and in the Ignore condition, it was *MS E* = 0.037(95%*CI*: 0.026 − 0.047). This suggests that people were able to accurately maintain a representation of the outcome experiences, even when instructed to ignore certain outcomes to make their choices. The estimates were similar in the discrete condition in Experiment 2 [Control: 0.029(95%*CI*: 0.026 − 0.031); Ignore: 0.025(95%*CI*: 0.021 − 0.029]; however, they slightly reduced, suggesting that people found the relative judgment format easier. When participants saw continuous outcomes, the MSE of their frequency estimates increased (Control: 0.134(95%*CI*: 0.125 − 0.143; Ignore: 0.125(95%*CI*: 0.113 − 0.137), suggesting that people found it more difficult to represent the continuous outcome experiences accurately.

Linear regression was used to predict the squared error in frequency estimates from the key experimental manipulations (Instruction [ignore or Control], Instruction Timing [Before or After], Distribution Type [Discrete or Continuous], Numeracy [High or Low]). All variables and their two-way interactions were entered into the model. We evaluated the importance of predictors in terms of the delta in explained variance (R-squared) between a model with the predictor and a model without. The results are displayed in Figure 3; three predictors emerged as significant. First, people who were presented with outcomes from a continuous distribution made considerably larger errors than people who saw discrete outcomes (Δ = 0.93, *b* = 0.102[0.095, 0.109]). Second, people with high numeracy scores made smaller errors than people with low numeracy scores (Δ = 0.013, *b* = − 0.038[− 0.044, − 0.031]). Finally, the distribution type interacted with numeracy, with people with high numeracy scores showing a larger difference between the discrete and continuous distributions (Δ = 0.008, *b* = −0.061[−0.075, −0.048]).

In sum, the analysis of frequency judgments demonstrated strong memory representations, unaffected by either the ignore manipulation or the timing of instructions. This is noteworthy, given that frequency judgments were introduced unexpectedly, indicating that people form explicit memory representations by default. However, the significant disparity between errors in discrete and continuous tasks also highlights the limitations of these memory representations, particularly among individuals with lower numerical abilities..

## Discussion

What is the nature of memory representations formed during an experience-based decision-making task? To experimentally find out, we designed a critical manipulation and asked people to ignore certain outcomes. Importantly, if people were able to ignore these target outcomes, the relative attractiveness of the risky option should increase, and their choice preference should change. Moreover, when the to-be-ignored outcomes are eliminated from the process of evaluation, the resulting decision problems reduce to ones that have been studied extensively in the decisions-from-experience literature. This, in turn, allowed us to predict choice patterns based on extensive prior findings (Wulff et al., 2018)).

We found clear evidence that people were able to update their impressions, with participants responding to the “ignore” instruction showing an increased preference for the risky option. This pattern suggests that they had access to distinct memory representations of the sequentially experienced episodes at the time of evaluation and choice. As a result, they could adaptively adjust their reliance on these episodes based on the instruction to ignore, rather than obeying them in a rigid or automatic way. The shift in preference was not moderated by the timing of instructions or the complexity of the problem. It was, however, affected by people’s numeric abilities, suggesting a role of individual differences. In contrast to choice, responses in a surprise frequency judgment task were not affected by the ignore instruction and were accurate in low- but not high-complexity problems. Overall, these findings establish that individuals do construct explicit memory representations of experienced episodes and rely on them when making their choices.

### Implications for models of memory and decision-making

#### Standard RL: No episodic memory

Our findings challenge traditional reinforcement learning (RL) models. They assume incremental value updating without the storage of individual episodes. As a consequence, standard RL cannot explain the flexible revision of preferences in response to the ignore instructions, highlighting the need to include memory-based mechanisms in modeling choice.

#### Episodic representations with limited flexibility

Several models in the experience-based decision-making literature, including MINERVA-DM (Dougherty et al., 1999), Decision by Sampling (Stewart, Chater, & Brown, 2006), MEM-EX (Hotaling et al., 2019), instance-based learning (Gonzalez & Dutt, 2011), and simple episodic RL models (Bornstein et al., 2017; Gershman & Daw, 2017), explicitly store individual episodes and use them to guide choice, often producing effects of recency or frequency. These models are highly similar in principle: they maintain memory of past outcomes but rely on fixed retrieval and weighting rules, limiting flexibility. While they can predict choice behavior in many standard experience-based tasks and produce reasonably accurate frequency judgments for discrete distributions, they are less well-suited to continuous outcome environments. The increased memory demands of continuous outcomes and the need for flexible adjustment in response to task instructions, as observed in response to the present ignore instruction, highlight the limitations of these fixed-retrieval models.

#### Episodic Deep RL: Flexible episodic control

A central implication of our findings is that they are consistent with a recent class of reinforcement learning models known as episodic controllers in Deep RL (Blundell et al., 2016; Botvinick, Wang, Dabney, Miller, & Kurth-Nelson, 2020). These models extend traditional RL by combining access to specific past experiences with the ability to generalize across them. Our results suggest that human decision-makers may operate in a similar way: they preserved distinct representations of individual episodes but used them flexibly, adjusting how these episodes influenced their choices specifically in response to the task instructions. This pattern cannot be fully captured by standard RL or by rigid DfE / simple episodic RL models; instead, it points to a system that integrates detailed episodic access with flexible, instruction-sensitive control over choices.

People’s memory representations were also robust across discrete and continuous distribution types, but this robustness was evident for choice rather than for frequency estimation. The finding that participants could revise their preferences in continuous-outcome environments while producing considerably worse frequency judgments is especially informative about the nature of memory representations. It suggests that participants are able to retain noisy information within the declarative memory system without relying on integrative strategies, allowing them to flexibly adjust choices in response to task instructions without maintaining fully precise or detailed episodic records of every outcome. One possibility is that participants encode continuous outcomes in categories or higher-level groupings, which facilitates adaptive decision-making but undermines accurate frequency recall. More broadly, these findings suggest that human memory may strike a compromise: retaining sufficient detail to enable flexible, episodic-like control over choices, while compressing or restructuring information in ways that favor functionality over fidelity (Markant & Gureckis, 2014; Mason et al., 2022)).

## Limitations

Our study cannot precisely identify the nature of memory representations in decisions from experience. First, our design only assessed memory access at the time of choice via the ignore instruction. While this reveals that participants can flexibly retrieve and use episodic information, it does not provide insight into how outcomes were encoded—for example, whether participants spontaneously categorized outcomes or integrated them differently during learning (see (**?**) on the role of memory at encoding). In addition, memory was measured indirectly through choice behavior and surprise frequency judgments. This, in turn, limits the precision with which we can characterize episodic representations.

Second, the paradigm uses a relatively small set of numeric outcomes. This constrains our ability to determine the granularity or exact form of memory representations (e.g., precise values versus categorical or approximate encoding). Third, our paradigm relies on monetary lotteries, which imply numerical information and a specific context (i.e., money, budget, finance). This context may elicit different processes than others involving more complex options in other domains (e.g., health, environmental, or social decisions; (Olschewski et al., 2023)). Finally, the design precludes decisions about sampling termination; in real-world settings, deciding when to stop sampling may influence how memory representations are formed and used (Hogarth & Einhorn, 1992; Wulff et al., 2018; **?**). Future work varying sample size, outcome precision, task domain, and including richer memory measures will be necessary to address these limitations.

## Conclusion

The current studies underscore key considerations for models of experience-based decision-making. People retain episodic memories of past outcomes, which they can flexibly access to adjust preferences according to task instructions. Individual differences, like numeric ability, also impact this process, highlighting the need for models that incorporate both flexible memory retrieval and individual variation. Recent models, such as episodic controllers in Deep RL, offer ways to integrate detailed memory with flexible use. However, the broader challenge is understanding how humans balance memory precision with functional flexibility. This includes compressing or categorizing outcomes to allow adaptive decision-making, even when exact outcome frequencies are not maintained.

## Declarations

### Research Transparency Statement

#### General Disclosures

Conflicts of Interest: The authors declare no conflicts of interest. Funding: This research was not supported by any specific grant. Artificial Intelligence: Generative AI tools (specifically, ChatGPT and Grammarly) were used solely to assist with language refinement and minor editorial improvements. They were not used for writing, data analysis, interpretation, or the generation of original content. Ethics: This research received approval from the Max Planck Institute for Human Development’s internal ethics board.

#### Study 1 and 2

Preregistration: No aspects of the studies were preregistered. Materials: Study instructions for both experiments and the experimental software for Study 1 are publicly available (https://osf.io/d7u5e). Data: All primary data are publicly available (https://osf.io/d7u5e). Analysis Scripts: All analysis scripts are publicly available (https://osf.io/d7u5e).

## Author contributions

Conceptualization: A.L., M.L., R.H., D.U.W.; Methodology: A.M., M.L., D.U.W; Formal Analysis: A.M., D.U.W. Resources: M.L., D.U.W. Writing – Original Draft: A.M., D.U.W. Writing – Review & Editing: A.M., M.L., R.H., D.U.W. Visualization: D.U.W. Funding Acquisition: R.H.

## References

Blundell, C., Uria, B., Pritzel, A., Li, Y., Ruderman, A., Leibo, J. Z., … Hassabis, D. (2016). Model-Free Episodic Control. arXiv. doi: 10.48550/arxiv.1606.04460

Bornstein, A. M., Khaw, M. W., Shohamy, D., & Daw, N. D. (2017). Reminders of past choices bias decisions for reward in humans. Nature Communications, 8(1), 15958. doi: 10.1038/ncomms15958

Botvinick, M., Wang, J. X., Dabney, W., Miller, K. J., & Kurth-Nelson, Z. (2020). Deep Reinforcement Learning and Its Neuroscientific Implications. Neuron, 107(4), 603–616. doi: 10.1016/j.neuron.2020.06.014

Christiano, P. F., Leike, J., Brown, T., Martic, M., Legg, S., & Amodei, D. (2017). Deep reinforcement learning from human preferences. Advances in neural information processing systems, 30. Retrieved from.neurips.cc/paper_files/paper/2017/file/d5e2c0adad503c91f91df240d0cd4e49-Paper.pdf},

Cokely, E. T., Galesic, M., Schulz, E., Ghazal, S., & Garcia-Retamero, R. (2012). Measuring risk literacy: The berlin numeracy test. Judgment and Decision making, 7(1), 25–47. doi: 10.1017/S1930297500001819

Daw, N. D., & Doya, K. (2006). The computational neurobiology of learning and reward. Current Opinion in Neurobiology, 16, 199–204. doi: 10.1016/j.conb.2006.03.006

Decker, J. H., Otto, A. R., Daw, N. D., & Hartley, C. A. (2016). From Creatures of Habit to Goal-Directed Learners: Tracking the Developmental Emergence of Model-Based Reinforcement Learning. Psychological Science, 27(6), 848–858. doi: 10.1177/0956797616639301

Dougherty, M. R. P., Gettys, C. F., & Ogden, E. E. (1999). MINERVA-DM: A memory processes model for judgments of likelihood. Psychological Review, 106(1), 180–209. doi: 10.1037/0033-295X.106.1.180

Erev, I., Glozman, I., & Hertwig, R. (2008). What impacts the impact of rare events. Journal of Risk and Uncertainty, 36(2), 153–177. doi: 10.1007/s11166-008-9035-z

Gershman, S. J., & Daw, N. D. (2017, January). Reinforcement Learning and Episodic Memory in Humans and Animals: An Integrative Framework. Annual Review of Psychology, 68(1), 101–128. doi: 10.1146/annurev-psych-122414-033625

Gonzalez, C., & Dutt, V. (2011). Instance-based learning: Integrating sampling and repeated decisions from experience. Psychological Review, 118(4), 523–551. doi: 10.1037/a0024558

Hertwig, R., Barron, G., Weber, E. U., & Erev, I. (2004). Decisions from experience and the effect of rare events in risky choice. Psychological science, 15(8), 534–9. doi: 10.1111/j.0956-7976.2004.00715.x

Hertwig, R., & Wulff, D. U. (2022). A description–experience framework of the psychology of risk. Perspectives on psychological science, 17(3), 631–651. doi: 10.1177/1745691621102689

Hogarth, R. M., & Einhorn, H. J. (1992). Order effects in belief updating: The belief-adjustment model. Cognitive psychology, 24(1), 1–55. doi: 10.1016/0010-0285(92)90002-J

Hotaling, J., Jarvstad, A., Donkin, C., & Ben, N. (2019). How to Change the Weight of Rare Events in Decisions from Experience. Psychological Science. doi: 10.1177/0956797619884324

Lengyel, M., & Dayan, P. (2007). Hippocampal Contributions to Control: The Third Way. Advances in Neural Information Processing Systems, 20, 889–896.

Lindskog, M., Winman, A., & Juslin, P. (2013). Calculate or wait: Is man an eager or a lazy intuitive statistician? Journal of Cognitive Psychology, 25(8), 994–1014. doi: 10.1080/20445911.2013.841170

Markant, D. B., & Gureckis, T. M. (2014). Is it better to select or to receive? Learning via active and passive hypothesis testing. Journal of Experimental Psychology: General, 143(1), 94– 122. doi: 10.1037/a0032108

Mason, A., Madan, C. R., Simonsen, N., Spetch, M. L., & Ludvig, E. A. (2022). Biased confabulation in risky choice. Cognition, 229, 105245. doi: 10.1016/j.cognition.2022.105245

Olschewski, S., Luckman, A., Mason, A., Ludvig, E. A., & Konstantinidis, E. (2023). The Future of Decisions From Experience: Connecting Real-World Decision Problems to Cognitive Processes. Perspectives on Psychological Science. doi: 10.1177/17456916231179138

Palminteri, S., & Lebreton, M. (2022). The computational roots of positivity and confirmation biases in reinforcement learning. Trends in cognitive sciences, 26(7), 607–621. doi: 10.1016/j.tics.2022.04.005ExternalLink

Plonsky, O., Teodorescu, K., & Erev, I. (2015). Reliance on small samples, the wavy recency effect, and similarity-based learning. Psychological Review, 122(4), 621–647. doi: 10.1037/a0039413

Silver, D., Hubert, T., Schrittwieser, J., Antonoglou, I., Lai, M., Guez, A., … others (2017). Mastering chess and shogi by self-play with a general reinforcement learning algorithm. arXiv. doi: 10.48550/arXiv.1712.01815

Spektor, M. S., & Wulff, D. U. (2021). Myopia drives reckless behavior in response to over-taxation. Judgment and Decision Making, 16(1), 114–130. doi: 10.1017/S1930297500008329

Spektor, M. S., & Wulff, D. U. (2024). Predecisional information search adaptively reduces three types of uncertainty. Proceedings of the National Academy of Sciences, 121(47), e2311714121. doi: 10.1073/pnas.2311714121

Stewart, N., Chater, N., & Brown, G. D. (2006). Decision by sampling. Cognitive Psychology, 53(1), 1–26. doi: 10.1016/j.cogpsych.2005.10.003

Sutton, R. S., & Barto, A. G. (1998). Reinforcement learning: An introduction. Cambridge, MA: MIT Press.

Weber, L. A., Yee, D. M., Small, D. M., & Petzschner, F. H. (2025). The interoceptive origin of reinforcement learning. Trends in Cognitive Sciences. doi: 10.1016/j.tics.2025.05.008

Wulff, D. U., Mergenthaler-Canseco, M., & Hertwig, R. (2018). A meta-analytic review of two modes of learning and the description-experience gap. Psychological Bulletin, 144(2), 140–176. doi: 10.1037/bul0000115

Wulff, D. U., & Pachur, T. (2016). Modeling valuations from experience: A comment on Ashby and Rakow (2014). Journal of Experimental Psychology: Learning, Memory, and Cognition. doi: 10.1037/xlm0000165

